# CRANBERRY: A Coarse-grained RNA Model with Sugar Puckering and Noncanonical Base Pairing

**DOI:** 10.64898/2026.01.12.699131

**Authors:** Yiheng Wu, Riccardo Alessandri, Aria E. Coraor, Xiangda Peng, Pablo F. Zubieta Rico, Korbinian Liebl, Cedric Salamé, Kha Trinh, Tobin R. Sosnick, Juan J. de Pablo

## Abstract

We introduce a new coarse-grained model “CRANBERRY” that incorporates sugar puckering and non-canonical base pairing, two factors central to RNA structure and dynamics, yet rarely included in most coarse-grained models. Our model is parameterized through a contrastive divergence approach, combined with fine-tuning strategies to improve accuracy in generating disordered states, a feature that is critical for the accurate description of thermodynamics. This two-stage training procedure greatly enhances cooperative folding behavior. Due to these advances, CRANBERRY maintains native-like structures with RMSD distributions comparable in scale to those obtained with all-atom force fields. Furthermore, CRANBERRY exhibits better agreement with experimental data on stacking free energies and disordered structures measured by Small Angle X-ray Scattering. In addition, CRANBERRY undergoes repeated folding and unfolding transitions of the ggcGCAAgcc tetraloop, reaching a minimum RMSD of 1.4 Å from an initially extended strand. CRANBERRY predicts melting temperatures in agreement with experimental values, and with a greater cooperativity than all-atom predictions.

## Introduction

The structure of RNA affects a wide range of processes, from biological function to therapeutic targeting, providing information on gene (1), protein translation (2, 3), and catalysis (4, 5). The static perspective of the RNA structure has evolved into an ensemble-based view of RNA dynamics (6–11), which likely is critical for cellular functions (1, 11) and human diseases (12). RNAs exhibit rugged folding free energy landscapes (FES) (7, 13–16). This opens up the possibility that the biologically relevant conformations are not always their ground states. These observations help motivate the development of accurate and efficient models for sampling RNA structural ensembles and thermodynamics. Our aim is not to compare our model directly with models designed primarily for structure refinement or prediction, such as deep-learning models (17–21), or certain Monte Carlo (MC) models (22–26). These models are effective for structure prediction, but generally not well suited for modeling RNA folding transitions and/or thermodynamics (27–29).

Several all-atom force fields for RNA have been developed (30, 31). A widely used model is OL3 (32), which offers good performance near the native basins of various RNAs. However, biologically relevant RNA motions on the microsecond time scale and beyond are difficult to capture with all-atom models due to computational limitations (30, 33), even for systems as small as tetraloops (34–39). Additionally, RNA force fields (31, 40–44) such as OL3 (32) or DES-AMBER (33) face challenges in correctly predicting disordered states and thermodynamics while maintaining the stabilities of the native structure. This apparent trade-off among native states, disordered states, and thermodynamics suggests an underlying imbalance between various energy terms (36, 43).

To accelerate computation while systematically addressing the imbalance issue, we choose to develop a coarse-grained (CG) model that accelerates sampling of the RNA ensembles and thermodynamics by reducing the number of degrees of freedom and smoothing free energy landscapes. (30, 45) The increased speed also makes it practical to train and refine the model iteratively. Within CG RNA models, two parameterization strategies dominate (30, 45–47). Knowledge-based (KB) models derive statistical potentials from structural databases via reference-state formalisms (48). BRiQ (23, 24), SPQR (25, 26), SimRNA (22), isRNA (49, 50), RACER (51–53), and HiRE-RNA (54, 55) are examples of this type. Reference-state methods face a self-consistency problem and require iterative parameterizations, which are computationally expensive (49, 56). Most CG models of this type aim only to predict or refine structures (30, 45), with notable exceptions such as RACER (51–53) and Hi-RE RNA (54, 55) that can fold RNA motifs *de novo*. However, thermodynamic quantities can be difficult to define rigorously.

The second approach - models based on thermodynamics - usually starts with nearest-neighbor (NN) parameters (57) and aims to match melting temperatures and other thermodynamic properties. oxRNA (58), TIS (5, 59, 60), Martini (61), and 3SPN.2 (62) belong to this category. They capture thermodynamics well, but typically require prior knowledge of RNA structures, rarely produce *de novo* folding transitions, and finally often assume A-form helices with limited support for noncanonical base pairs (6, 7). More recently, iConRNA combined an intermediate-resolution representation with effective explicit Mg^2+^ ions and reproduced the conformational properties of poly(rA) and poly(rU), folding and melting of small structured RNAs, and sequence-, temperature-, and ion-dependent RNA phase behavior (63).

RACER, originally knowledge-based, was subsequently optimized against unfolding free energies, but this thermodynamic optimization reduced native stabilization, echoing the trade-offs observed in all-atom force fields (43). We use RACER as the primary CG benchmark because it provides published results for the three properties examined here: native-state stability, disordered-state ensembles, and folding thermodynamics.

In summary, KB and thermodynamics-based models face four main challenges: 1) costly/ill-posed reference states (KB); 2) limited noncanonical pairing (thermodynamics); 3) difficulty achieving both thermodynamics and folding; 4) underrepresented sugar-puckering dynamics that influence binding, catalysis, and primer extension (64–67). (SPQR represents sugar puckering, but by discrete MC moves.)

To address these challenges, we develop CRANBERRY (**C**oarse-grained **R**NA model with **AN**isotropic **B**as**E** interactions and **R**ibose **R**ing puckering d**Y**namics): 1) We use the contrastive divergence (ConDiv) method (68–71), which has proven effective for parameterizing the CG protein model *Upside* (72). This trajectory-based iterative method does not require a postulated reference state and is computationally less expensive than rigorous reference state approaches. 2) We further optimize by fine-tuning stacking free energies and disordered structures. This two-step training is analogous to the dual-target training used to improve *Upside* (73). 3) We design energy functions for non-canonical base pairings that avoid degeneracy and have a clear correspondence to all-atom base-pairing geometries. 4) We develop a many-body bonded potential to represent the collective transition between the two dominant sugar-puckering states. CRANBERRY strives to provide a favorable balance among native-state stability, disordered-RNA dimensions, and folding thermodynamics across the benchmarks examined here. CRANBERRY outperforms the CG model RACER in these three benchmarks. Additionally, CRANBERRY predicts melting temperatures within 15 K of their experimental values and shows greater cooperativities (sharper melting curves) than all-atom force fields. Moreover, CRANBERRY can fold the tetraloop ggcGCAAgcc into native conformations *de novo*, reversibly, with an RMSD_min_ of 1.4 Å. CRANBERRY also reproduces the destabilization of the C2’-endo state in an A-form helix.

## Materials and Methods

### Model

In CRANBERRY, each nucleotide is represented by 6 sites: one for the phosphate group, two for the sugar, and three for the base (Fig. 1A). Compared to isRNA2 (50) or RACER (51–53), we use two sites rather than one, C3’ (S3) and C2’ (S2), to represent the sugar. These two sites allow us to model the sugar pucker transition between C3’-endo and C2’-endo, as discussed in the section Sugar Puckering Energy Function. The base mapping is the same as in RACER (51–53) (Fig. 1C). We note that the mapping for the base does not have a significant impact on the model energy function as long as three sites are chosen, because the base stacking and base pairing interactions only take the center of geometry and the normal vector of the bases as input (Section Materials and Methods: Nonbonded interactions). Following RACER, placing the sites near the base edges better captures the excluded volume of the bases. We put a virtual site at each base’s center of geometry, termed BC (Base Center), and another virtual site, termed BN (Base Normal), such that the vector from BC to BN defines the base-plane normal (Fig. 1B). The normal vector is chosen to point towards the 5’ direction in a canonical A-form helix. Both BC and BN are used in base stacking and pairing potentials to define base-plane orientation. Additionally, BC is used in steric calculations to better represent the excluded volume.

**Fig. 1.**
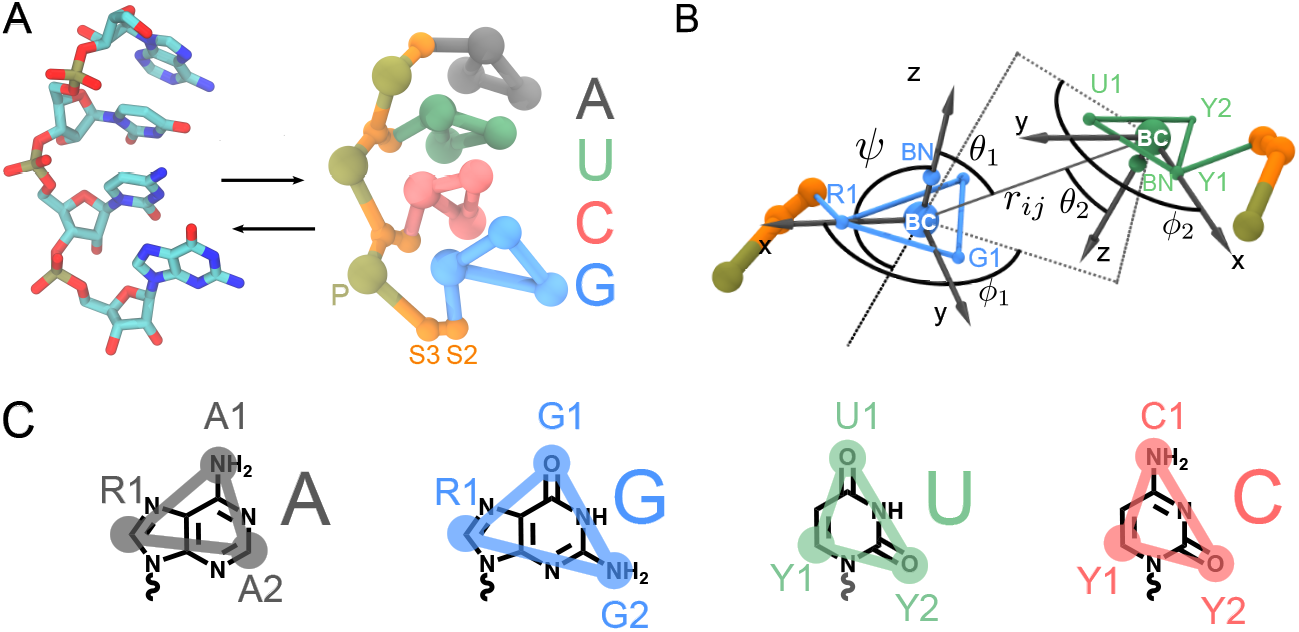
The CRANBERRY model. (A) Mapping between all-atom and coarse-grained resolutions, with coarse-grained sites placed on the phosphate, C3’ and C2’ of the sugar, and 3 atoms of the base (B) Positions of virtual sites BC (base center) and BN (base normal), and definitions of degrees of freedom *r*_*ij*_, *θ*_1_, *θ*_2_, *ϕ*_1_, *ϕ*_2_, *ψ* between two interacting bases (G-U in the figure). BC is placed at the base center of the geometry, and BN is positioned so that the vector from BC to BN defines the base normal. The direction is chosen such that BC-BN points towards the 5’ direction in an A-form helix. *r*_*ij*_ is the distance between base *i* and *j*. We define a local coordinate system for each base, aligning the z-axis with the base normal and the x-axis with the vector BC-R1/Y1. *θ*_1_, *θ*_2_ are angles between 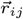 and the z-axis of each base. To calculate *ϕ*_1_, *ϕ*_2_, we first project the vector 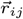 onto each base plane, then calculate the angle between the projected vector and the x-axis of each base. Note that *ϕ*_1_, *ϕ*_2_ range from [0, 2*π*]. *r*_*ij*_, *θ*_1_, *ϕ*_1_ are the coordinates of BC of base *j* in the local spherical coordinate system centered at BC of base *i*. (Mathematical definitions can be found at Eq. 8) (C) Mapping of the four base types, overlaid with their chemical structures.

### Data sets

To parameterize the model, we curated a fragment and a ConDiv data set. The fragment data set was used to parameterize bonded potentials and provide initial parameters for base stacking and pairing potentials. The ConDiv data set was used to train base stacking, pairing, and spline potentials via ConDiv (68–71).

The fragment data set was constructed by filtering the non-redundant 3D RNA structure data sets (74), removing structures with a resolution higher than 3 Å. This data set includes the RNA components of the RNA-protein and RNA-ligand complexes (e.g., ribosomes and riboswitches). This data set was then used to compute various statistics: 1) coarse-grained degrees of freedom (bonds, angles, and dihedral angles) and phase angles (via the Cremer and Pople method (75)), used to parameterize the linear regression model for predicting sugar puckering (Section Materials and Methods: Sugar puckering energy function); 2) histograms of bonds, angles, and dihedrals used for parameterizing bonded potentials; 3) histogram of degrees of freedom of base pairing (*r, θ*_1_, *θ*_2_, *ϕ*_1_, *ϕ*_2_, *ψ* (Fig. 1B)), used to create initial parameters for base pairing before ConDiv training. Base pairs are identified by DSSR (76). This data set contains 1150 RNAs, 127,293 nucleotides, and 162,333 valid base pairs. It is usually used for local features and should be regarded as a fragment library. As a result, we did not filter chain breaks or perform MD simulations on this data set.

The ConDiv data set was constructed by further filtering the non-redundant list using the NAKB (Nucleic Acid Knowledgebase) webserver (77). This data set includes RNA-only structures (crystal structures, Cryo-EM, or NMR) composed exclusively of standard nucleotides. The molecular weight range is restricted to 6.6-33 kDa, which corresponds to ~ 20-100 nucleotides. This range was chosen to ensure that the trajectories remain computationally feasible during training. We further excluded quadruplexes (which require modeling explicit base-ion interactions) and structures containing ligands. We also removed any structure that contains chain breaks. In addition, we applied a 3 Å resolution cutoff for X-ray and Cryo-EM structures. In total, this data set contained 242 NMR structures, 24 X-ray structures, and one fluorescence-derived model.

### Bonded energy functions

The bonded potential, *U*_b_, includes bond (*U*_bond_), angle (*U*_angle_), and dihedral (*U*_dihedral_). The bond and angle terms are harmonic. The dihedral term uses the combined bending-torsion potential (CBT) (78), which mitigates numerical instabilities when three atoms become colinear and the dihedral becomes degenerate. Several approaches exist to address dihedral instabilities (78–80); we selected CBT for its simplicity.

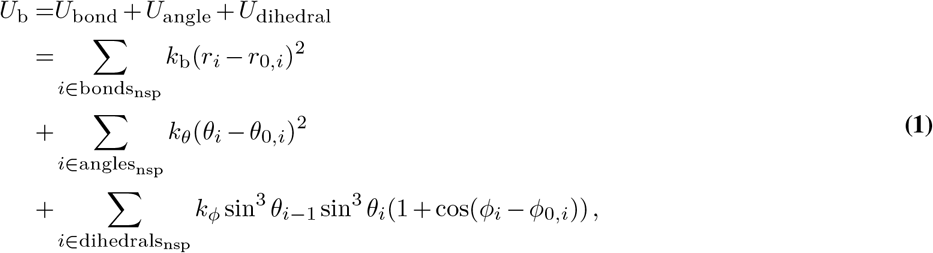

where *θ*_*i*−1_, *θ*_*i*_ are the angles formed by 3 of the 4 particles corresponding to the dihedrals (angle 1-2-3 or 2-3-4 when the dihedral is formed by 1-2-3-4), and the factors sin^3^ *θ*_*i*_ are included to prevent diverging forces as internal angles approach 180° (78). Bonds, angles, and dihedrals not related to sugar puckering are denoted bonds_nsp_, angles_nsp_, dihedrals_nsp_ (where “nsp” stands for “not sugar pucker”). The parameters are listed in Supplementary Table S1-3.

### Sugar puckering energy function

Sugar puckering refers to the out-of-plane motion of the five-membered ribose ring (81– 85). This collective motion couples the degrees of freedom of the backbone to the base (86, 87). RNA ribose most frequently adopts two conformations: C3’-endo and C2’-endo. This motion is well described by the phase angle (75, 85, 88), a collective variable derived from the coordinates of the five ring atoms. The Cremer-Pople definition (75) of the phase angle projects all five ribose sites to the mean plane of the ribose (Fig. 2B), and then calculates the phase angle from the projection distance *z*_*j*_ for each site *j*

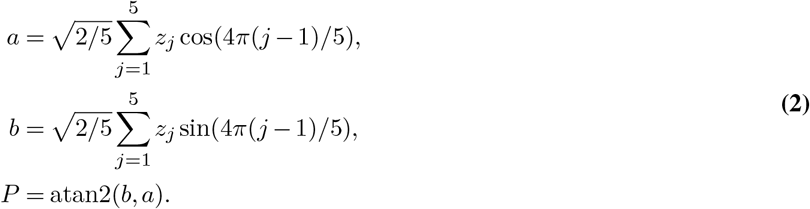

**Fig. 2.**
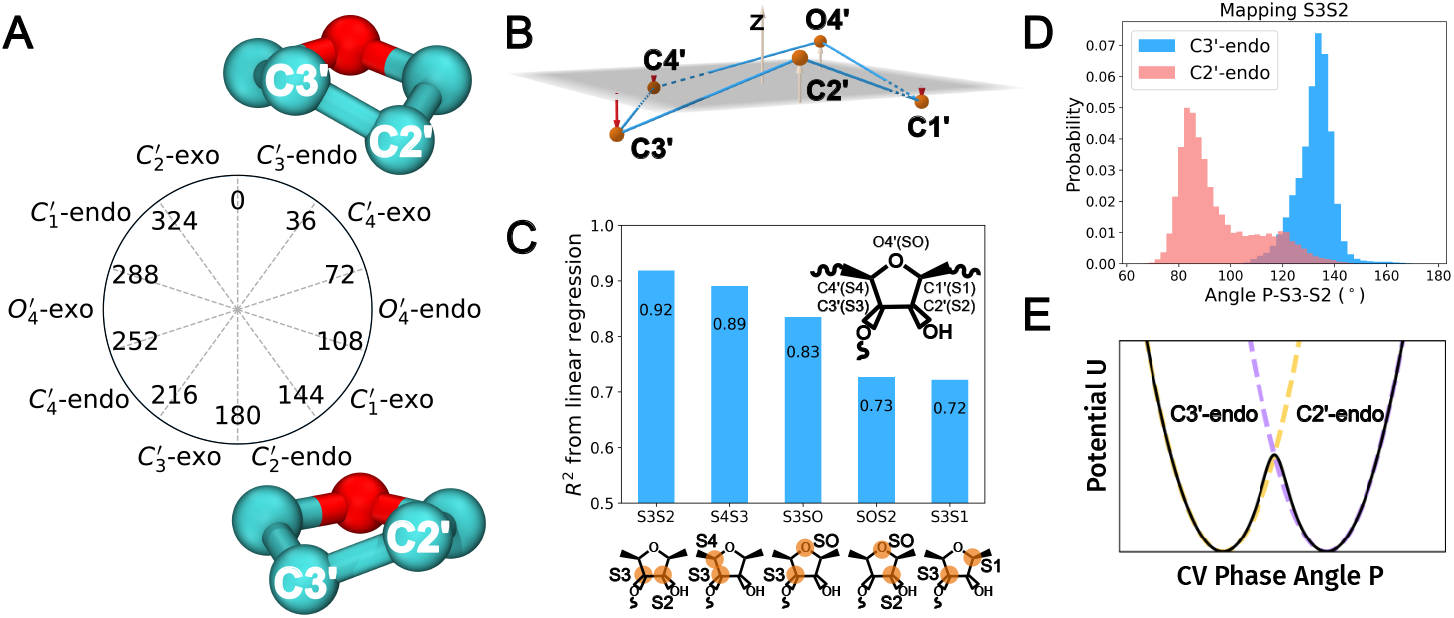
Schematics and results for sugar puckering energy function design. (A) Possible conformations of ribose and the corresponding phase angles, with C3’-endo and C2’-endo visualized. (B) Schematics for the Cremer–Pople method for calculating the phase angle. The coordinates of each atom in the ribose are projected to the mean plane, and the length of the projection is used to calculate the phase angle. (C) *R*^2^ of the test set from the linear regression if different mapping schemes are chosen. (D) Example distribution of coarse-grained angle P-S3-S2 with respect to sugar puckering states. (E) A schematic representation of the sugar puckering bonded energy function, which can be thought of as a collective variable that provides smooth interpolation between C3’-endo energy functions and C2’-endo energy functions.

The phase angle can distinguish all possible conformations of the ribose, including C3’-endo (phase angle in the range 0–36°) from C2’-endo (157°–180°) (Fig. 2A). Similarly, north and south conformations are defined as conformations in the north (0° *< P <* 90° or 270° *< P <* 360°) or the south (90° *< P <* 270°) part of the phase angle plot (89). However, the phase angle calculation requires all five ring atom coordinates. This raises the question of how to model this collective transition and distinguish between C3’-endo and C2’-endo in a coarse-grained model. We address this by developing a linear regression model to predict the phase angle from the coarse-grained sites and use it to interpolate between two sets of bonded energy functions corresponding to the C3’-endo and C2’-endo states (Fig. 2E).

The predicted coarse-grained phase angle *P*_*j*_ is defined by

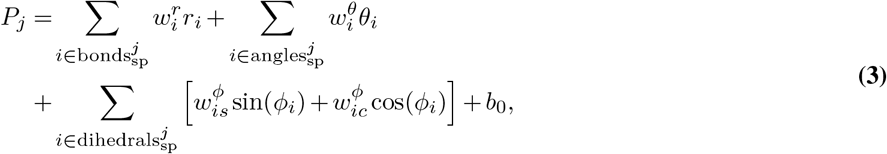

which is a linear combination of coarse-grained bonds, angles, and the sine and cosine of dihedrals. We denote the bonds, angles, and dihedrals that are highly correlated with sugar pucker (for example, coarse-grained angle P-S3-S2, Fig. 2D) as 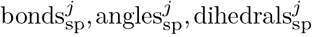 with *j* denoting the nucleotide index, and “sp” indicating the relevance of sugar puckering.

We first compared candidate two-site sugar mappings using the complete set of 15 descriptors (coarse-grained bonds, angles and dihedrals) available for each mapping. The optimized mapping, with S3 and S2 placed at C3’ and C2’, respectively, achieved a test *R*^2^ of 0.92 (Fig. 2C), indicating that these sites effectively distinguish C3’-endo from C2’-endo conformations. After selecting the C3’–C2’ (S3–S2) mapping, we constructed the production sugar-puckering collective variable using a reduced set of 10 descriptors. This reduced model achieved a test *R*^2^ of 0.90, and its coefficients, which were incorporated into the force field, are reported in Supplementary Table S7. We used the sine and cosine of the dihedral angles as input features, rather than the dihedral angle itself, to ensure that the predicted phase angle varies smoothly when a dihedral angle crosses the periodic boundary between 180° and −180°. The parameters defining the predicted phase angle *P*_*j*_ are listed in Supplementary Table S7. We use the predicted phase angle *P*_*j*_ to interpolate between bonded energy functions parameterized separately for C3’-endo and C2’-endo (Fig. 2E), as defined in Eq. 4.

The sugar puckering contribution to the energy function, *U*_pucker_, is then represented by a many-body bonded potential. This many-body bonded potential interpolates between bonded potentials parameterized for C3’-endo *U*_3_ and C2’-endo *U*_2_. *U*_3_ and *U*_2_ are standard bond, angle, and dihedral potentials, using the same formulas as *U*_bond_, *U*_angle_, *U*_dihedral_ in Eq. 1. The difference lies in the sets of bonds, angles, and dihedrals to which they are applied. This choice is because some coarse-grained degrees of freedom strongly correlate with sugar puckering states (Fig. 2D). The parameters for these bond/angle/dihedrals are listed in Supplementary Table S4-6.

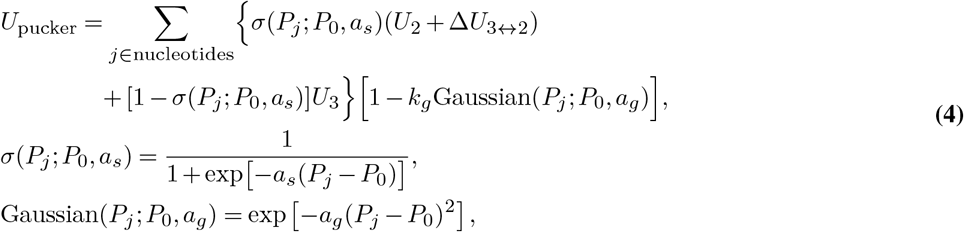

where *σ*(*P*_*j*_, *P*_0_, *a*_*s*_) serves as a switching function that transitions the bonded potential between the C3’-endo and the C2’-endo. Gaussian(*P*_*j*_, *P*_0_, *a*_*g*_) controls the energy barrier between C3’-endo and C2’-endo. *P*_*j*_ is the predicted phase angle for the sugar in the nucleotide *j* as defined in Eq. 3, where *P*_0_ = 90° is the switching threshold, and *a*_*s*_ = (0.085°^−1^), *a*_*g*_ = (4.38 × 10^−6^ °^−2^) control the sharpness of the switching function and the Gaussian functions, respectively (Fig. 2E). Δ*U*_3↔2_ = 130 kJ mol^−1^ is parameterized by the population ratio for free nucleotides. *k*_g_ = 0.947 controls the overall strength of the Gaussian modification part. This sugar puckering potential is applied to each nucleotide independently. For simplicity, if the 5’-phosphate is missing, the terminal nucleotide is modeled by the bonded terms as in C3’-endo.

### Nonbonded interactions

The nonbonded potential, *U*_nb_, includes excluded volume interactions (*U*_wca_), base stacking (*U*_stacking_), base pairing (*U*_pairing_), electrostatics (*U*_elec_), and spline-based interactions between backbone (phosphate P and sugar sites S2 and S3) and base (*U*_spline_).

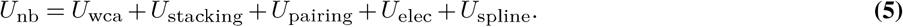

The excluded volume interactions are represented by the Weeks–Chandler–Andersen (WCA) potential (90).

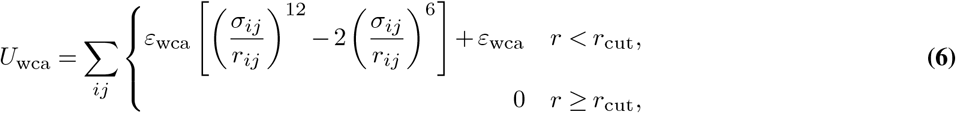

where the energy parameter *ε*_wca_ is globally set to be 5 kJ mol^−1^, and *σ*_*ij*_ = (*σ*_*i*_ + *σ*_*j*_)*/*2 is the average diameter of the site. *r*_*ij*_ denotes the distance between particles. The *σ*_*i*_ parameters are listed in Supplementary Table S8. Note that, in addition to the standard sites, the BC virtual site also participates in WCA interactions.

The base stacking potential is anisotropic and involves one radial switching function and three angular switching functions between two bases.

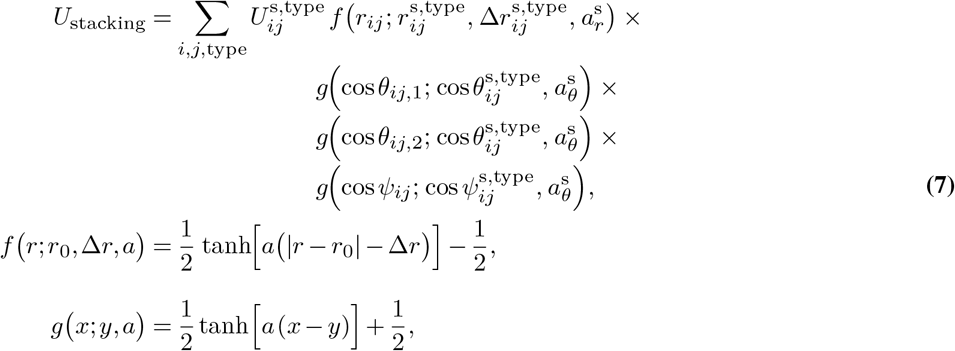

where *i* and *j* are the indices of the nucleotides. Visual representations of *r*_*ij*_, *θ*_1_, and *θ*_2_ are shown in Fig. 1B, where cos 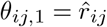, cos 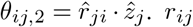 is the distance between BC of the nucleotide *i* and that of nucleotide *j*, and 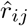 is its unit vector. 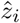 and 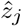 are unit vectors along the z-axes of bases *i* and *j*, respectively. Possible types are 33, 55, and 35, corresponding to interactions via 3’-faces and 5’-faces. The 5’-face is defined as the side of the base plane containing the atom C5’. In canonical A-form helices, neighboring bases stack via 3’-5’ faces (35), and this type of stacking is termed ‘normally stacked’. Accordingly, 33 and 55 types of stacking are denoted as ‘reversely stacked’ (Fig. 6B). All the parameters are provided in Supplementary Table S9. The experimental enthalpies (91) are asymmetric with respect to the swapping the first and second nucleotide type. For example, the enthalpy of UG is different from that of GU. Thus, we allow the interaction strength matrix for the 35 stacking 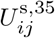 to be asymmetric. In contrast, the matrices for the 33 stacking 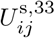 and 55 stacking 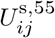 are symmetric by definition.

All three edges of a base can participate in base pairings, which are termed Watson–Crick (WC) edge, Hoogsteen edge, and sugar edge (92–94) (Fig. 3A). To differentiate different edges, we propose to use the *ϕ*_1_ and *ϕ*_2_ angles, which are the angles between the center of geometry vectors projected to the base plane and the x-axis of the local coordinate systems (Fig. 1B, defined in Eq 8). In a canonical GC WC–WC base pairing, *ϕ*_1_ and *ϕ*_2_ are both around 180° (Fig. 3B). In a sheared GA pair (Sugar–Hoogsteen), *ϕ*_1_ ≈ 290° and *ϕ*_2_ ≈ 60° (Fig. 3C). As indicated by these two examples, *ϕ*_1_ and *ϕ*_2_ are good indicators of base pairing geometry (Fig. 3A, *ϕ*_1_ and *ϕ*_2_ for all the base pairs in the fragment data set (Materials and Methods: Data sets), colored by the accurate assignment of base geometry performed using DSSR (76, 95, 96)). Different geometries are clearly well-separated in the scatter plot *ϕ*_1_ − *ϕ*_2_. Our definition of *r, θ*_1_, *θ*_2_, *ϕ*_1_, *ϕ*_2_, *ψ* is equivalent to that of the SPQR model (25). In the SPQR model (25), the 6D joint distribution of two bases (*r, θ*_1_, *θ*_2_, *ϕ*_1_, *ϕ*_2_, *ψ*) is simplified by assuming that the base orientations are independent given a fixed inter-base distance. In contrast, we retain the key correlation between the base orientations between *ϕ*_1_ and *ϕ*_2_ by decomposing different base pair geometries. This decomposition is required because of the different underlying all-atom hydrogen-bonding patterns. Taking into account base types, edge types, and base orientations, there are a total of 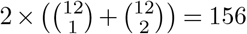 distinct base pairing geometries. We model only the 25 most frequent base pairing geometries found in the fragment data set, accounting for 87% of the base pairs, and leave the rest rare base pairings as zero energies.

**Fig. 3.**
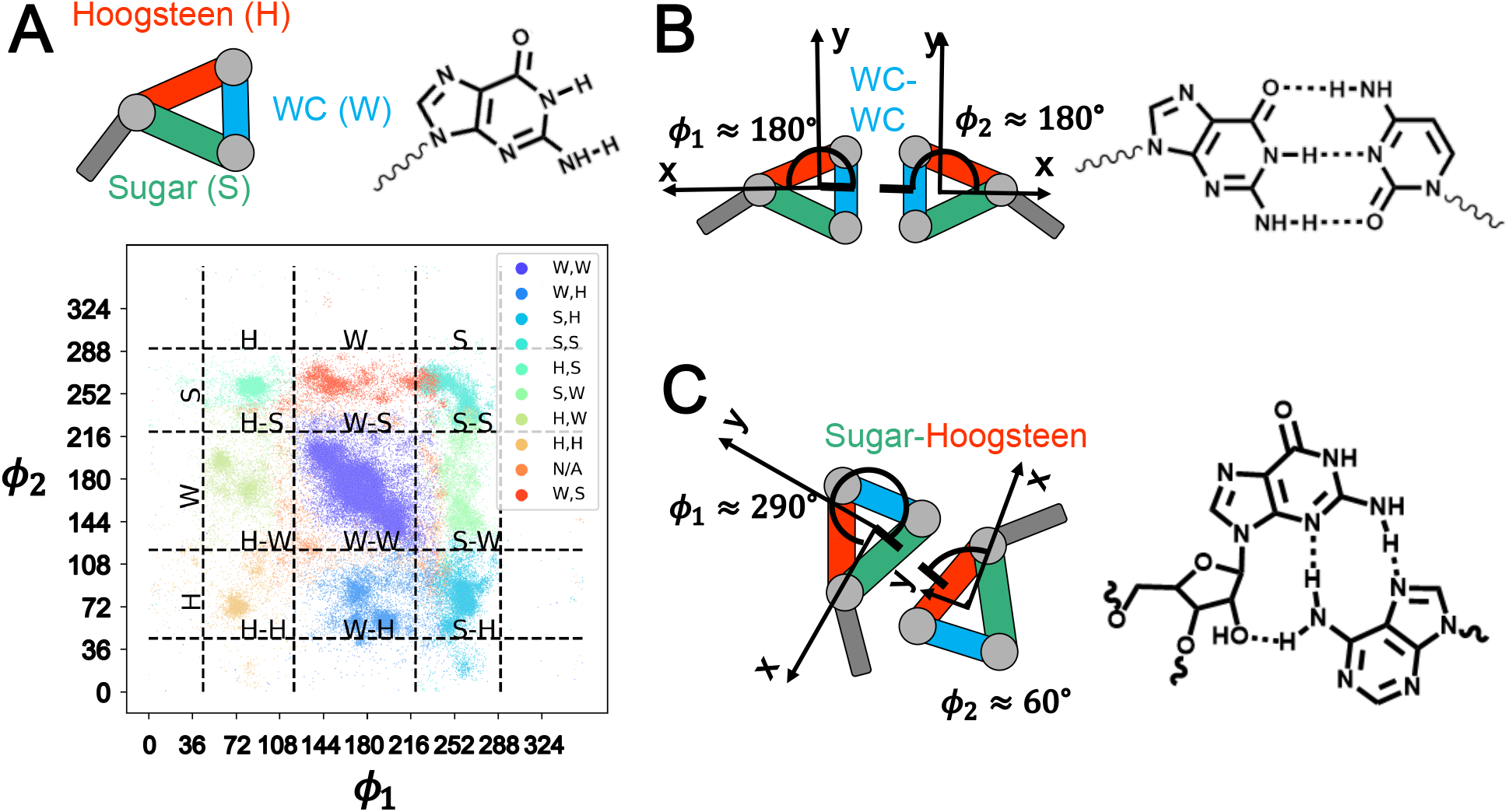
Schematic representations of base interaction edges and *ϕ*_1_-*ϕ*_2_ plot of base pairing. (A) Schematic of the three base interaction edges (Watson-Crick, Hoogsteen, and Sugar) (92–94) and the chemical structure of Guanosine. The *ϕ*_1_-*ϕ*_2_ (defined in Fig. 1B and Eq.8) plot illustrates a clear separation between different base geometries. The colors of the scatters are determined by DSSR (76). (B) Example of a WC–WC interaction in a GC pair, where both *ϕ*_1_ and *ϕ*_2_ are near 180°, along with the corresponding chemical structure. (C) Example of a Sugar–Hoogsteen interaction, where *ϕ*_1_ ≈ 290° and *ϕ*_2_ ≈ 60°, shown with its corresponding chemical structure

The base pairing potential is then modeled as an anisotropic interaction with one radial component and five angular components.

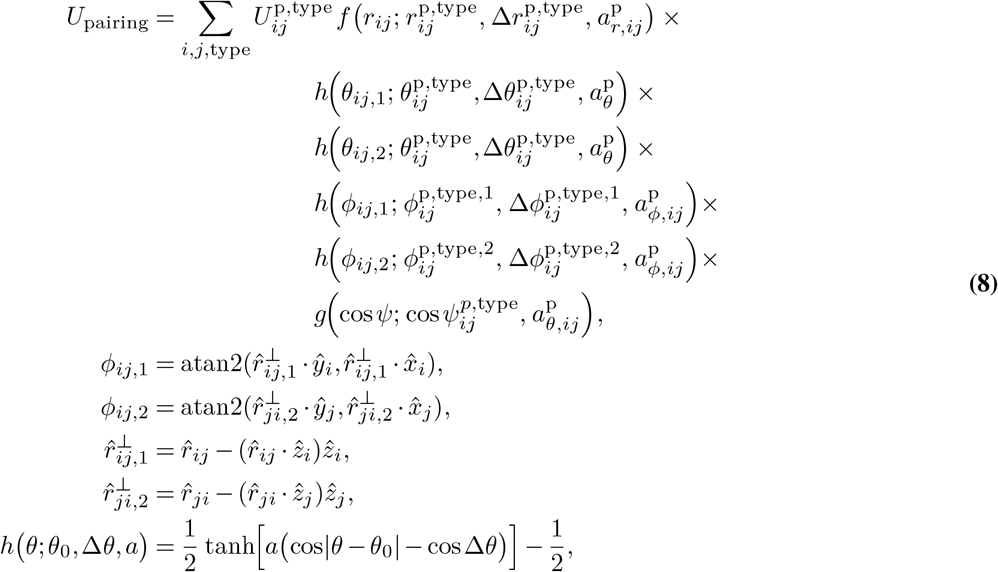

where 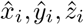 are the unit vectors along the local x/y/z-axis of each base (Fig. 1). *r*_*ij*_, *θ*_*ij*,1_, *ϕ*_*ij*,1_ are the BC coordinates of base *j* in the local spherical coordinate system of base *i*, with 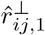 being the projection of 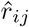 onto the plane of base *i* (Fig. 1B). Their frequencies are shown in Supplementary Fig. S1. These 25 geometries are denoted by “type” in Eq. 8. The corresponding parameters are tabulated in Supplementary Table S10. Base pairing between adjacent bases on the same strand is explicitly disallowed.

Electrostatic interactions are approximated using a Debye-Hückel type screened potential to account for salt effects, following the 3SPN.2 CG model for DNA (62)

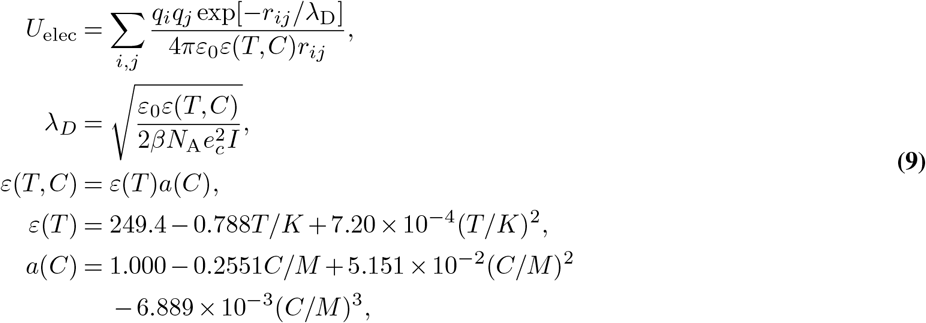

where *q*_*i*_ and *q*_*j*_ are the charges on phosphate *i* and *j, r*_*ij*_ is the inter-site distance, *λ*_*D*_ is the Debye screening length, *ε*_0_ is the dielectric permittivity of vacuum, *ε*(*T, C*) is the dielectric permittivity of the solution. *β* is the inverse thermal energy (*k*_B_*T*)^−1^, *N*_A_ is Avogadro’s number, *e*_*c*_ is the elementary charge, and *I* is the ionic strength of the solution. The dielectric permittivity of the solution *ε*(*T, C*) is a function of temperature *T* and the concentration of the NaCl solution *C*. For each simulated NaCl concentration *C*, we set *I* = *C*, as appropriate for a 1:1 electrolyte, and recalculate both *ε*(*T, C*) and *λ*_D_(*T, C*). Thus, each NaCl concentration is represented by a distinct screening length; no explicit ions are included, and all other force-field parameters are held fixed. Following 3SPN.2 (62), the phosphate charge is fixed at *q*_P_ = −0.6*e* at all concentrations to approximate the effect of counterion condensation (97, 98).

The spline potential provides a flexible way to represent the complex interactions of phosphate–phosphate, sugar–sugar, phosphate–sugar, phosphate–base (99), and sugar–base:

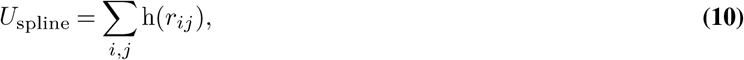

where *h*(*r*) is represented by natural cubic splines. For each pair of residue types, different spline functions are trained (Supplementary Table S11 and Supplementary Fig. S2). The spline functions between base sites are set to zero, since their interactions are already captured by base-stacking and base-pairing.

For all the nonbonded interactions, pairs of sites separated by three or fewer bonds are excluded. CRANBERRY includes approximately 1000 parameters in total.

### Langevin dynamics

Following 3SPN.2 (62), Langevin dynamics is used to propagate the dynamics of the system with a time step of 5 fs. To assign the damping constant, the Einstein relation is used

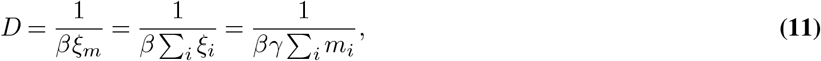

where *ξ*_*m*_ is the molecular friction coefficient, and *ξ*_*m*_ = ∑_*i*_ *ξ*_*i*_ where *ξ*_*i*_ is the friction coefficient of each site, and *i* is summing all the sites in the molecule. We ignore the difference in the damping coefficients between sites with different masses. We use an empirical scaling relation (100) to account for RNA length in diffusivity *D* = 4.58 × 10^−10^*N* ^−0.39^m^2^s^−1^, where N is the number of nucleotides. The elimination of microscopic degrees of freedom generally produces non-Markovian projected dynamics characterized by memory-dependent friction and correlated fluctuating forces (101, 102). Conventional coarse-grained models replace these effects with simplified Markovian equations and often smooth the free-energy landscape, thereby altering and typically accelerating dynamical timescales (103, 104). Consequently, our diffusion-based damping coefficient constrains the approximate center-of-mass diffusivity but does not establish the physical timescale of internal conformational relaxation or folding. Quantitative kinetic interpretation would require calibration against experimental rate or relaxation measurements, as demonstrated for 3SPN.2 DNA simulations (105, 106). These items should be considered when interpreting CRANBERRY’S time scale.

### Parametrization approach

Using the fragment data set, we manually optimized the bonded potential to correctly capture the data set’s distribution of bonded degrees of freedom (Supplementary Fig. S3). Then, we determine the steric repulsion parameters by two criteria. First, we compared the peak positions of the radial distribution function of all sites in all-atom simulations of an A-form helix in water with the sum of LJ diameter (2.8 Å) for the water and the diameter of the coarse-grained sites. Secondly, we ensure that for the canonical A-form helix 3ND4, the steric repulsion is minimal. Afterward, we initialize the base pairing and base stacking interactions with parameters derived from the fragment data set. We use all-zero values as initial parameters for spline interactions, since they are minor contributions to the total energy of the system. Following the 3SPN.2 parameterization protocols (62), the strength parameters for the stacking are initialized by scaling the experimental enthalpies (91).

After obtaining the initial parameters, we applied ConDiv training (71) to 235 entries in the 267-entry collection, with 32 static entries held out for validation. Experimental method and training role were treated separately. Of the 242 NMR-derived entries, 203 were used as multi-model training ensembles, whereas the remaining 39 were represented by a single fixed structure and assigned to the static training (20 entries) or static validation (19 entries) sets. Together with 24 X-ray structures and one fluorescence-derived model, the collection therefore comprised 32 static training entries, 32 static validation entries, and 203 multi-model NMR training entries. We approximate the reference distribution of these RNAs at room temperature via restrained simulations near their structures. Restrained simulations are performed by randomly selecting 3*N*_sites_ native contacts and then applying harmonic restraints to these contacts (71, 73). At each epoch, we run RNA trajectories in the training set for around 1 ns, and we calculate approximate derivatives of the Kullback-Leibler divergence between the reference distribution and the distribution produced by CRANBERRY (71). For each NMR entry and training epoch, three distinct deposited models were selected randomly without replacement and used to initialize three independent approximately 1-ns free simulations. Their parameter derivatives were averaged when calculating the CONDIV update. The 20 NMR-derived entries in the static training set were instead represented by a single fixed structure and were not subjected to three-model sampling. The Adam optimizer (107) is used for optimization. The ConDiv method is applied only to the nonbonded base pairing, stacking, and spline interactions, while the bonded terms are inferred from the fragment data set. We run 30 epochs to achieve convergence. We further fine-tune the stacking parameters by running benchmark simulations of di-nucleotide monophosphate for which the experimental stacking free energies are available (91). In this fine-tuning step, we only tune the strength parameter of the stacking potential 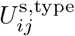, for the 35, 55, and 33 classes of stacking geometries. We found that the experimental stacking free energies Δ*G*_stack_ do not distinguish between normally and reversely stacked conformations. As a result, we use a hybrid scheme for this matching process, where we simultaneously match the experimental stacking free energies Δ*G*_stack_ (Supplementary Eq. 2) and the free energy difference between normally and reversely stacked Δ*G*_ns*/*rs_ (defined in Supplementary Eq. 3) estimated from OL3. Due to limited data, we assume that the strength of the 55 types to be the same as for the 33 types. The matching process is carried out by performing the secant method (108). Furthermore, we globally scaled the spline potential (backbone-backbone and backbone-side chain interactions) so that the Rg of rU30 matches the experimental value at 200 mM salt. We note that due to the limit of the training set size and information about dynamical ensembles, the ConDiv method tends to overfit, such that the structures might be overstabilized. This behavior is similar to the overstabilization tendency of all-atom force fields when the main goal is to stabilize different known DNA/RNA structures (31–33, 40, 109–111). Similar behavior is also observed for all-atom force fields (112) and coarse-grained models (71, 73) developed for proteins. A similar issue has recently been resolved by introducing regularization terms for protein binding affinities (113). Thus, this underscores the importance of including disordered RNA and thermodynamic fine-tuning, because they also serve as physical regularizers for the model.

### MD simulation protocols and FES calculations

The simulations of all-atom and coarse-grained force fields follow common practices (33, 40, 41, 53), with details of simulation setup and calculations of stacking, native fraction, and SAXS profile included in the Supplementary Methods.

## Results and Discussion

Having defined and parameterized CRANBERRY, we next evaluated its performance and compared it against OL3, DES-AMBER, and RACER. The models were tested for their ability to fold or maintain native structures, describe disordered states, and predict stacking free energies. We further tested CRANBERRY’s ability to reproduce the melting temperatures and to reversibly and *de novo* fold the ggcGCAAgcc tetraloop, and to predict sugar puckering FES for one nucleotide or a double helix.

### Structural stability

CRANBERRY generally stabilizes the motifs (tetraloops 1ZIH, 2KOC, double helix 3ND4, and pseudoknot 1L2X) better than RACER (Fig. 4 and Supplementary Fig. S4). For RACER, the pseudoknot 1L2X melts at 300 K, whereas CRANBERRY maintains its structure with an all-sites RMSD= 4–6 Å of the native structure. CRANBERRY can stabilize the two tetraloops to an RMSD of 1.5–5 Å. For the A-form helix 3ND4, CRANBERRY produces a more accurate structure than RACER, which exhibits a broad peak from 2.5 Å to 10 Å. For the two tetraloops, RACER predicts native-like structures around 6 Å.

**Fig. 4.**
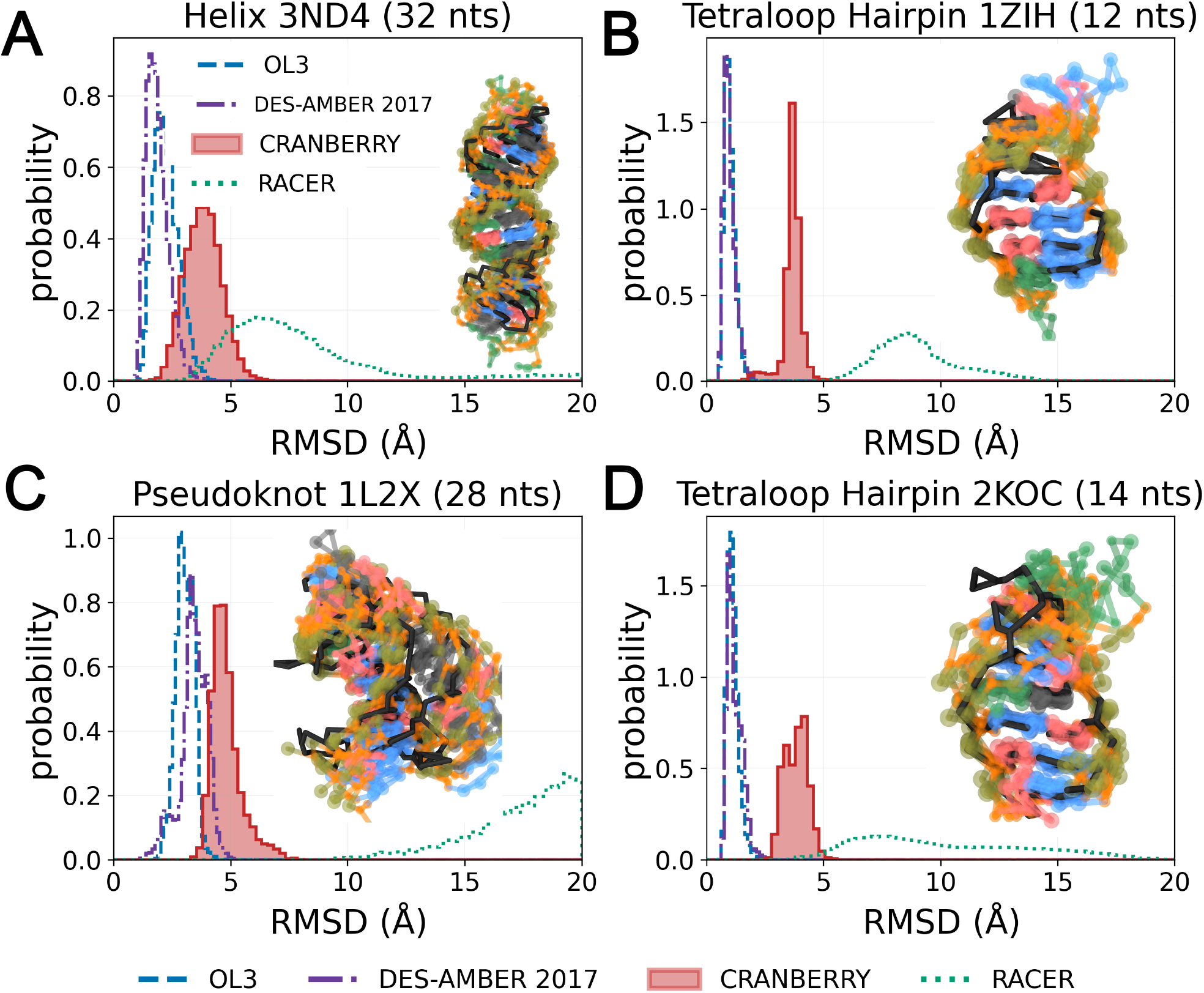
Native simulations for (A) a canonical A-form double helix 3ND4 (B) a tetraloop 1ZIH (C) a pseudoknot 1L2X (D) a tetraloop 2KOC by OL3(32), DES-AMBER 2017/TIP3P (33), CRANBERRY, and RACER (53). In each plot, the black structure is the crystal/NMR structure, and the colored transparent structures are the representative conformations of the CRANBERRY simulations (see Supplementary Methods for selection procedures).

The structural distributions of DES-AMBER and OL3 are largely concordant. For 1L2X, DES-AMBER yields slightly higher RMSDs than OL3, indicating that it samples conformations slightly farther from the experimental reference structures (33, 43). OL3 and DES-AMBER’s RMSDs values are generally 1–2 Å lower than CRANBERRY. Our slightly worse performance may have multiple origins. First, CRANBERRY does not model all base-pairing interactions and is coarse-grained, which makes it, in general, less accurate in capturing detailed interactions. However, there is also the possibility that OL3 and DES-AMBER overstabilize these native structures. This hypothesis is partially supported by OL3’s worse performance for disordered RNA, where it produces an overcollapsed state for rU30 (Fig. 5A) and rU40 (33) with a radius of gyration (Rg) 10 Å below experimental measurements (Fig. 5F). This interpretation is supported by later stacking free energies benchmarks as shown in Fig. 6, and that has been reported previously (40, 41). This suggests that OL3/DES-AMBER’s interactions might be too “sticky,” similar to what has been observed for all-atom protein force fields (112, 114, 115).

**Fig. 5.**
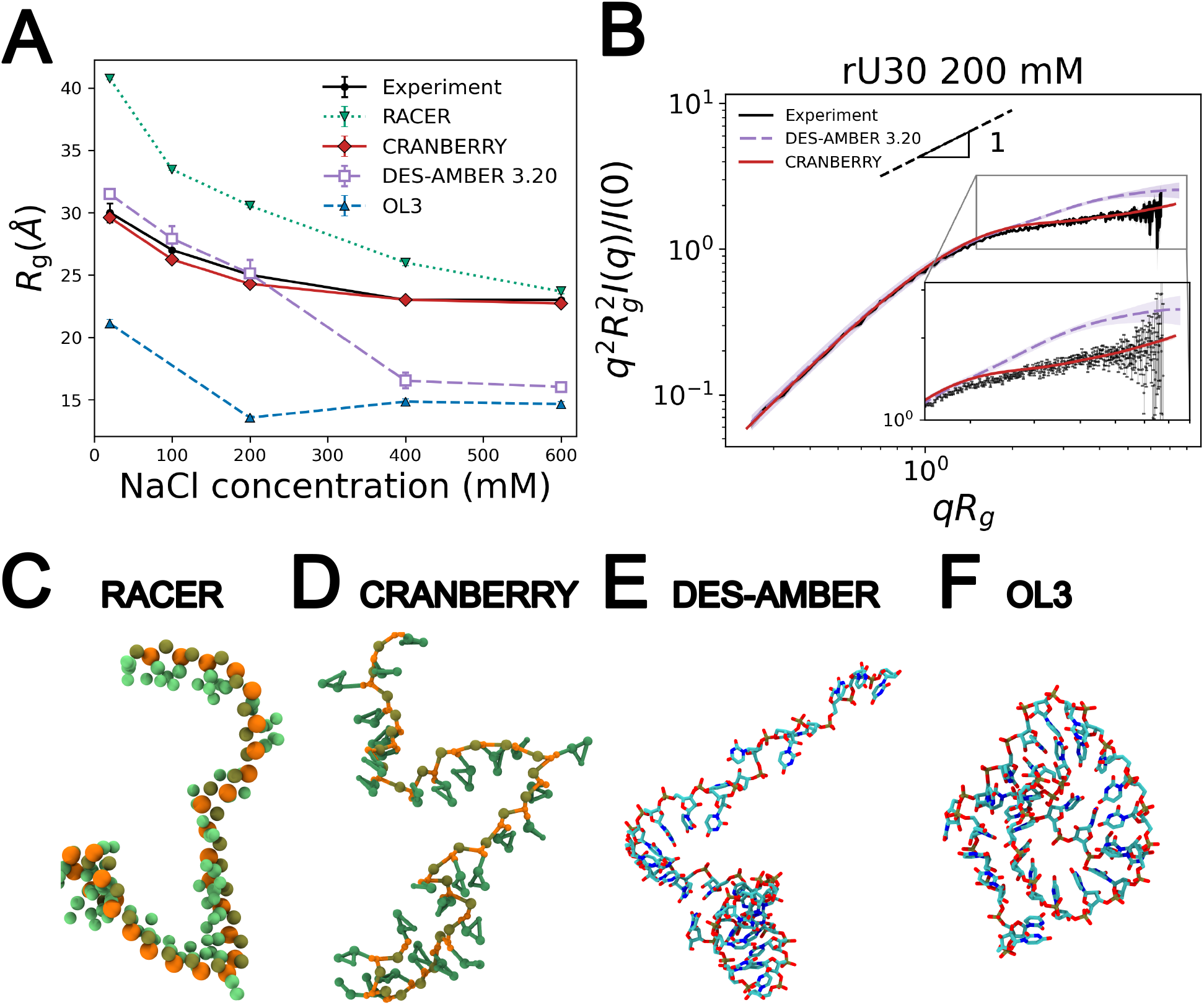
Radii of gyrations and scattering behaviors for disordered RNAs as sampled with different atomistic and coarse-grained force fields in comparison to experiments (A)*R*_g_ of rU30 varying salt conditions, comparison between experimental data (119) and various force fields. The reported *R*_g_ values are ensemble averages over the final three quarters of each trajectory; error bars represent standard error of the block means using the last 3/4 of the trajectory. The sizes of the error bars for RACER, CRANBERRY, and OL3 are close to the sizes of the markers. (B) Dimensionless Kratky plot of rU30 under 200 mM NaCl, showing comparison between experimental data, CRANBERRY, and DES-AMBER; asymptotic rod-limit scaling is indicated by a dashed black line. (C-F) Final configuration of rU30 under 200 mM NaCl using RACER(53), CRANBERRY, DES-AMBER/TIP4P-D (111) or OL3/TIP3P (32). (Note that here we switched to DES-AMBER 3.20 (111) rather than DES-AMBER 2017 (33) as 3.20 improves the predictions on the disordered RNA systems, thus serving as a better benchmark.)

**Fig. 6.**
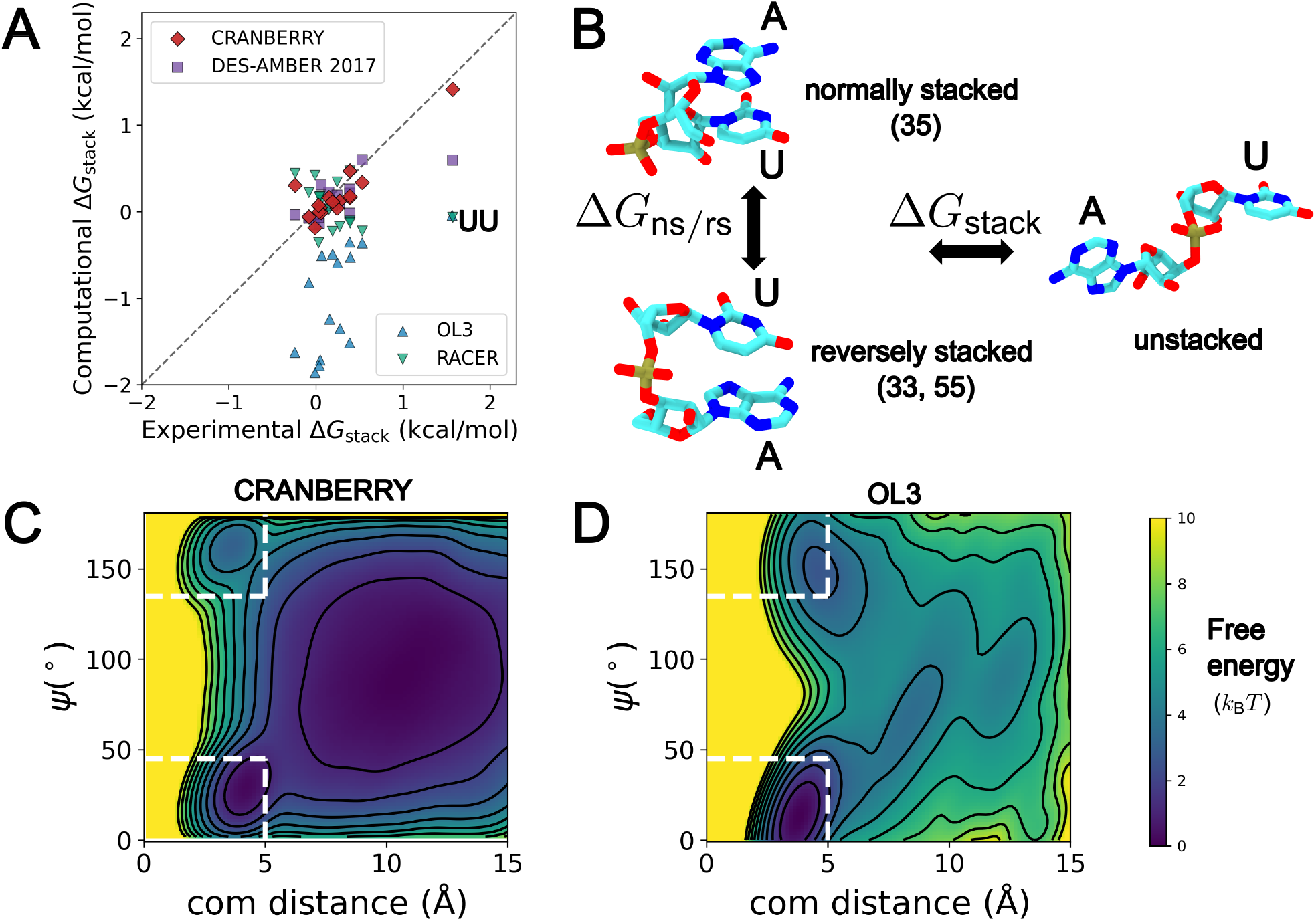
Stacking free energies. (A) Comparison between experimental (91) and computational Δ*G*_stack_. The scatter points with 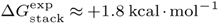 correspond to UU stacking. The DES-AMBER 2017 force field is used with TIP4P-D water models. (B) Snapshots of unstacked, normally stacked, and reversely stacked states (defined in the section Materials and Methods: Nonbonded interactions). (C-D) FES of dinucleotide monophosphate ApU, under CRANBERRY and OL3, respectively. The coordinates are the center-of-mass distance between two bases and the angle between the normal vectors of the two base planes. The contour lines are plotted every 0.25 *k*_B_*T*. The white rectangles denote the regimes of stacked conformations.

Because global RMSD does not directly report preservation of the native interaction network, we additionally quantified the level of native base pairs, including noncanonical interactions. In 1ZIH, the native G5–A8 G–A interaction (AH–GS) was populated by approximately 1% of the CRANBERRY ensemble, compared with 99% of the DES-AMBER ensemble, as estimated from the second half of each trajectory. These trajectory-based occupancies should not necessarily be interpreted as equilibrium populations. In particular, the near-unity DES-AMBER occupancy may partly reflect limited sampling from a native starting structure. Temperature-dependent folding simulations of a closely related 10-nt GCAA tetraloop reported an overall native-state population of approximately 50% near room temperature (33), which is a proxy to the occupancy of the individual G–A interaction for DES-AMBER.

The structurally more complex 1L2X pseudoknot contains seven native noncanonical base pairs identified by DSSR, four of which correspond to base-pairing geometries explicitly represented among the 25 geometries in CRANBERRY. The occupancies of these four interactions ranged from 0 to 1.73% with CRANBERRY, compared with 83.84–100% with OL3 and 22.46–100% with DES-AMBER. CRANBERRY’s directional base-pairing interactions were parameterized conservatively to reduce the risk of kinetically trapping overly stabilized contacts. Nevertheless, the low occupancies observed here suggest that the current interactions are underestimated for at least some native noncanonical geometries. Pair-specific experimental occupancies are not available, however, so these results do not establish whether the high occupancies obtained with the all-atom force fields are quantitatively accurate. The individual occupancies, DSSR classifications, and geometric criteria are provided in Supplementary Table S12, Supplementary Methods while more discussions can be found in Supplementary note 1.

### SAXS measurements of disordered RNAs

CRANBERRY’s simulations produce SAXS (Small Angle X-ray Scattering) data that are very similar to the experimental measurements on rU30 (Fig. 5A, B). CRANBERRY’s ability to reproduce native structures and unfolded conformations with a high degree of all-or-none cooperativity implies that our training procedure captures the delicate balance between various energy terms (interactions of the backbone side-chain, base stacking, electrostatic repulsions, sugar puckering bonded interactions, and internal stiffness of the chain).

All-atom methods produce a mixture of results. OL3 produces a collapsed conformation for the disordered rU30, with Rg ~ 10 Å below the experimental values (Fig. 5A, F). Similar overcompaction was previously reported for rU40 33. Pairing OL3 with the OPC water model can alleviate this excessive compaction (37, 116); however, recent work indicates that OPC can also destabilize certain RNA native structures (117).

DES-AMBER 3.20 shows closer agreement with experiment at low to intermediate salt concentrations. Its *R*_*g*_ values are slightly above the experimental values in 20 and 100 mM NaCl and close to experiment at 200 mM, but it samples overly compact conformations at 400 and 600 mM, thereby overestimating the salt-induced compaction. For this benchmark, we use DES-AMBER 3.20 (111), which updates the Lennard–Jones interaction between water and the nonbridging phosphate oxygens, instead of DES-AMBER 2017 (33). This revised phosphate–water interaction is particularly relevant for solvent-exposed disordered RNA ensembles. In contrast, DES-AMBER 2017 exhibits transient hairpin-like conformations (33) and, in rU30 simulations, produces *R*_*g*_ values approximately 4–5 Å below experiment (Supplementary Fig. S5). The DES-AMBER version used for each of the remaining benchmarks is specified separately in the Supplementary Methods. For RACER, the Rg value matches the experiment at high salt, but exhibits higher salt dependency compared to experiments at low to intermediate salt regimes (Fig. 5A).

At 200 mM NaCl, the CRANBERRY rU30 ensemble produces a scattering profile in excellent agreement with experiment (Fig. 5B). Although the overall spline strength was calibrated against *R*_g_ at this concentration, the scattering-profile shape was not a fitting target. CRANBERRY also reproduces the experimental profiles across the tested NaCl-concentration range (Supplementary Fig. S6). DES-AMBER exhibits slightly higher slopes in the high q range compared to the experiments, suggesting the simulated rU30 might be slightly too rigid at short length scales along its backbone. This rigidity may stem from the over-stabilization of UU stacking observed in DES-AMBER (Fig. 6A). CRANBERRY’s agreement with experiment is physically consistent with its explicit representation of sugar puckering, which previous SAXS and smFRET work identified as an important determinant of the contour length of poly(rU) (118).

We also examined rA30, for which SAXS/CD measurements reveal base stacking and ion-dependent conformational organization (119). CRANBERRY simulations sample two conformational populations, comprising relatively extended, locally stacked conformations and more compact conformations (Supplementary Figs. S8 and S10, with further discussions in the Supplementary note 3-4). The simulated ensemble is somewhat more compact than experiment in medium to high salts, indicating that CRANBERRY may have poorer effective solvent quality for rA30. Nevertheless, the overall range of simulated *R*_*g*_ values across the examined salt concentrations is comparable to that observed experimentally.

### Stacking free energy

As part of our parameterization procedure, CRANBERRY was fine-tuned against the experimental stacking free energies and reproduced both their overall scale and sequence variation (Fig. 6A, *R*^2^ = 0.79). Thus, this agreement reflects performance on a fitting target rather than an independent prediction. Among the comparison models, DES-AMBER with TIP4P-D provides the closest overall agreement with experiment (Fig. 6A, *R*^2^ = 0.56). OL3 exhibits a larger overestimation of ~ 1 kcal · mol^−1^, while RACER generally avoids overstabilization except for UU stacking. These results are broadly consistent with previous benchmarks (40, 41). However, the available experimental dataset is limited and several decades old; revisiting these measurements would provide a stronger parameterization target, and CRANBERRY can be refitted as improved data become available.

Three states emerge in the FES under the coordinates of the center of mass distance and angles between base normal vectors *ψ*: normally stacked, reversely stacked, and unstacked (Fig. 6C and D, and Supplementary Fig. S12). We note that the experimental stacking free energies are insufficient to fully parameterize the stacking across normally stacked (35), and reversely stacked (33, 55) orientations (Fig. 6B and the section Materials and Methods: Nonbonded interactions for definitions of 33, 35, and 55). Future experimental measurements distinguishing these stacked states, or more accurate all-atom force fields, could aid in parameterization of stacking interactions in the coarse-grained models.

### A. Melting and *de novo* folding of tetraloops and helices

Although transferability across wide temperature ranges is an inherent problem in coarse-graining (46, 47), CRANBERRY predicts the melting temperatures of ggcGCAAgcc and cacag within 15 K of the experimental values, comparable to the performance of DES-AMBER (33) (Fig. 7C and G). However, the cooperativity (sharpness of the melting curves) is steeper and closer to the experimental behavior of CRANBERRY (Fig. 7C and G).

**Fig. 7.**
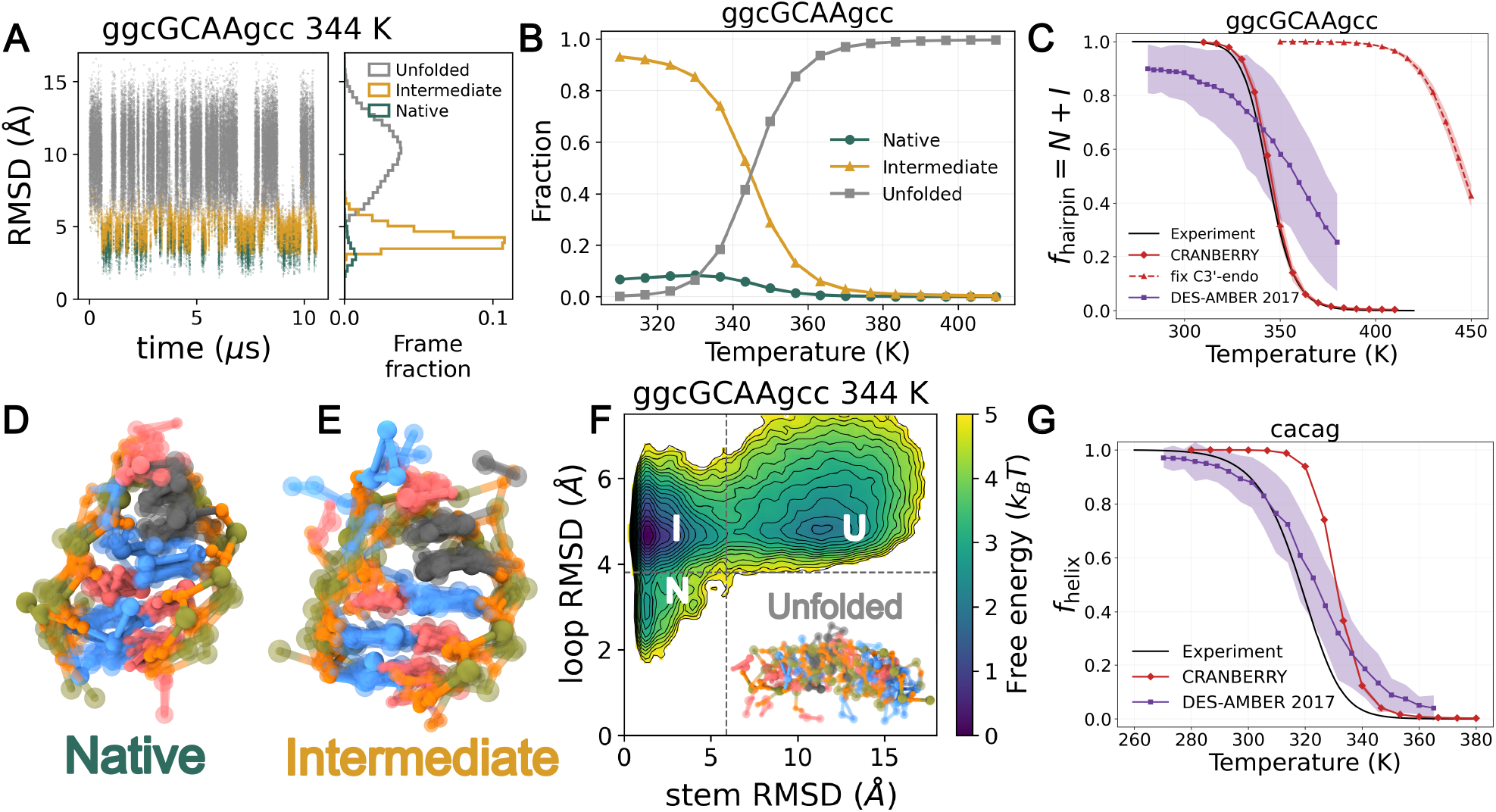
Melting and *de novo* folding of tetraloops and helices. (A) *de novo* folding of ggcGCAAgcc showing RMSD (against PDB: 1zih nt 2-11, frame 9 as the representative frame) over time at 344 K. The definition of native, intermediate, and unfolded can be found in Supplementary Methods. (B) The fraction of native, intermediate and unfolded conformations over temperature. (C) Melting curve of ggcGCAAgcc by plotting the fraction of hairpin conformations (*f*_hairpin_ = *f*_native_ + *f*_intermediate_), using DES-AMBER (taken from Ref. (33)) or CRANBERRY, with or without fixing sugar puckering states to C3’-endo. The experimental curve is plotted using the van ‘t Hoff equation (*K*_eq_ = exp(−Δ*H*^0^*/RT* + Δ*S*^0^*/R*)) and experimental folding enthalpy and entropy. For tetraloop folding *K*_eq_ = *f/*(1 − *f*), and for double-stranded cacag *K*_eq_ = 2*f/*[(1 − *f*)^2^(*c*_ss_*/c*^Θ^)] with the concentration for the single strands denoted as *c*_ss_ and the standard concentration *c*^Θ^ = 1 M. Errors are represented by the shaded area, and are calculated by blocking analysis. Because optical melting reports on the overall hairpin–coil transition rather than the native loop topology, the simulated observable is *f*_hairpin_ = *f*_native_ + *f*_intermediate_. (D,E) Representative conformations of native and intermediate states (selection procedures can be found in Supplementary Methods) produced by CRANBERRY, aligned by the stem. (F) FES of ggcGCAAgcc at 344 K with coordinates of RMSD of the stem (nt 1-3,8-10) and the loop (nt 4-7). The contour lines are plotted every 0.25 *k*_B_*T*. (G) Melting curves of the cacag double helix from experiments, DES-AMBER and CRANBERRY.

Near the melting temperature of ggcGCAAgcc, the FES contains three basins corresponding to native, intermediate, and un-folded conformations (Fig. 7F, state definitions are provided in the Supplementary Methods and indicated by the dashed lines). Starting from an extended strand, a 10-*µ*s CRANBERRY trajectory at 344 K samples all three states and undergoes repeated transitions among them, reaching a minimum RMSD of 1.4 Å (Fig. 7A). Because calibrating the internal kinetic timescale of coarse-grained models is challenging, the observed transition times should not be interpreted as physical folding rates.

The experimental melting curves were obtained by UV optical melting and analyzed as a two-state, unimolecular hairpin–coil transition (120). The absorbance change reports the ensemble-averaged hyperchromicity associated primarily with disruption of base stacking, but it does not distinguish the native GCAA loop topology or the G5–A8 noncanonical contact. Accordingly, within our three-state classification, the appropriate simulated observable is the total population of stem-folded conformations, *f*_hairpin_ = *f*_native_ +*f*_intermediate_, rather than *f*_native_ alone. Agreement with the optical melting curve therefore assesses overall hairpin formation, not the population of the fully native loop. At 300 K, the 4-*µ*s REMD ensemble contains approximately 94% intermediate, 5.8% native, and 0.08% unfolded conformations. Moreover, the native G5–A8 noncanonical base pair is formed in 0.8% of the ensemble (Supplementary Note 2). Thus, although CRANBERRY reproduces cooperative hairpin formation, the thermodynamic balance between the native and intermediate loop conformations remains imperfect.

The predicted melting of ggcGCAAgcc is more cooperative (Fig. 7C) than for DES-AMBER. A similar relationship between training against disordered conformations and folding cooperativity was observed during the development of *Upside* 1 and 2 for proteins (71, 73). *Upside* 1 exhibited weaker cooperativity when its energy function was trained only against native structures, whereas cooperativity improved in *Upside* 2 after native and disordered states were incorporated through a dual-target loss function (73). CRANBERRY’s fine-tuning approaches serve as a physical regularizer, providing an effect similar to the dual-target loss function for proteins, and producing cooperative folding behavior for CRANBERRY.

The less-cooperative behavior of DES-AMBER may be attributed to residual structures in the unfolded state arising from overstabilized nonbonded interactions in the all-atom force fields (112). Such overstabilization lowers the enthalpy of unfolded ensembles and consequently reduces the magnitude of the folding enthalpies. Indeed, all-atom models of proteins and RNAs, such as DES-AMBER, predict significantly lower folding enthalpies than experiments (33, 121), resulting in less cooperative melting transitions. Consistent with this tendency to overstabilize compact disordered conformations, DES-AMBER 2017 also underestimates the *R*_*g*_ of rU30 at high salt concentrations (Supplementary Fig. S7).

The simulated melting temperature of ggcGCAAgcc is increased by as much as ~ 100 K when all sugar puckering states are fixed to C3’-endo while keeping all other settings the same (Fig. 7C). This finding supports the hypothesis that sugar puckering exerts a non-negligible entropic effect on melting behavior. Recently, Zhang (122) proposed a back-of-the-envelope estimation of the entropy loss during the crystallization of polypropylene. The loss of entropy per dihedral is estimated to be *k*_B_ ln 3, because each dihedral is fixed to one of the three states when crystallization occurs. Applying analogous reasoning to ggcGCAAgcc, and assuming that each nucleotide samples both C3’-endo and C2’-endo in the unfolded ensemble but is restricted primarily to C3’-endo in the folded ensemble, gives an entropy contribution of approximately *k*_B_ ln 2 per nucleotide. Using experimental entropy and enthalpy values, the change in melting temperature caused purely by the fixing of sugar puckering is estimated to be *T*_*m*_ · 10*k*_B_ ln 2*/* |Δ*S*_exp_ | ≈ 60 K. This crude estimate agrees on the order of magnitude with our computational result, which underscores the importance of including sugar puckering in CG models to capture thermodynamics and melting behaviors accurately. This entropic interpretation also provides a direct physical connection to the widespread use of locked nucleic acids (LNAs) to enhance oligonucleotide thermal stability. The 2^′^-O,4^′^-C methylene bridge in an LNA nucleotide constrains its sugar to the C3’-endo/North conformation, thereby preorganizing the backbone for A-form pairing and reducing the conformational entropy lost upon folding or hybridization (123, 124). This preorganization contributes to the elevated melting temperatures of LNA-containing duplexes, although changes in stacking and hydration also contribute; the correspondence with our fixed-pucker calculation is therefore mechanistic rather than quantitative. Our argument on the sugar puckering entropy is also consistent with the theoretical analysis on the temperature transferability of coarse-grained models (47, 125), where including more fine-grained information provides better estimates of the entropy and hence achieves better melting behavior, although the lack of an explicit solvent makes the assignment of the melting entropy a challenge.

### Sugar puckering FES

CRANBERRY simulations of an isolated guanosine monophosphate (GMP) have near equilibrium between C3’-endo and C2’-endo at 300 K (Fig. 8A). CRANBERRY estimates that the difference in free energy between the C3’-endo and the C2’-endo for a guanosine monophosphate is Δ*G*_3↔2_ = *G*_C2’-endo_ − *G*_C3’-endo_ ≈ +0.2 *k*_B_*T* (C3’:C2’=55:45). In GMP NMR experiments, Δ*G*_3↔2_ is reported from −0.4 *k*_B_*T* (40:60) (89) to − 0.8 *k*_B_*T* (30:70) (82, 83). Thus, we consider CRANBERRY’s performance satisfactory for the thermodynamics of the C3’-endo:C2’-endo transition.

**Fig. 8.**
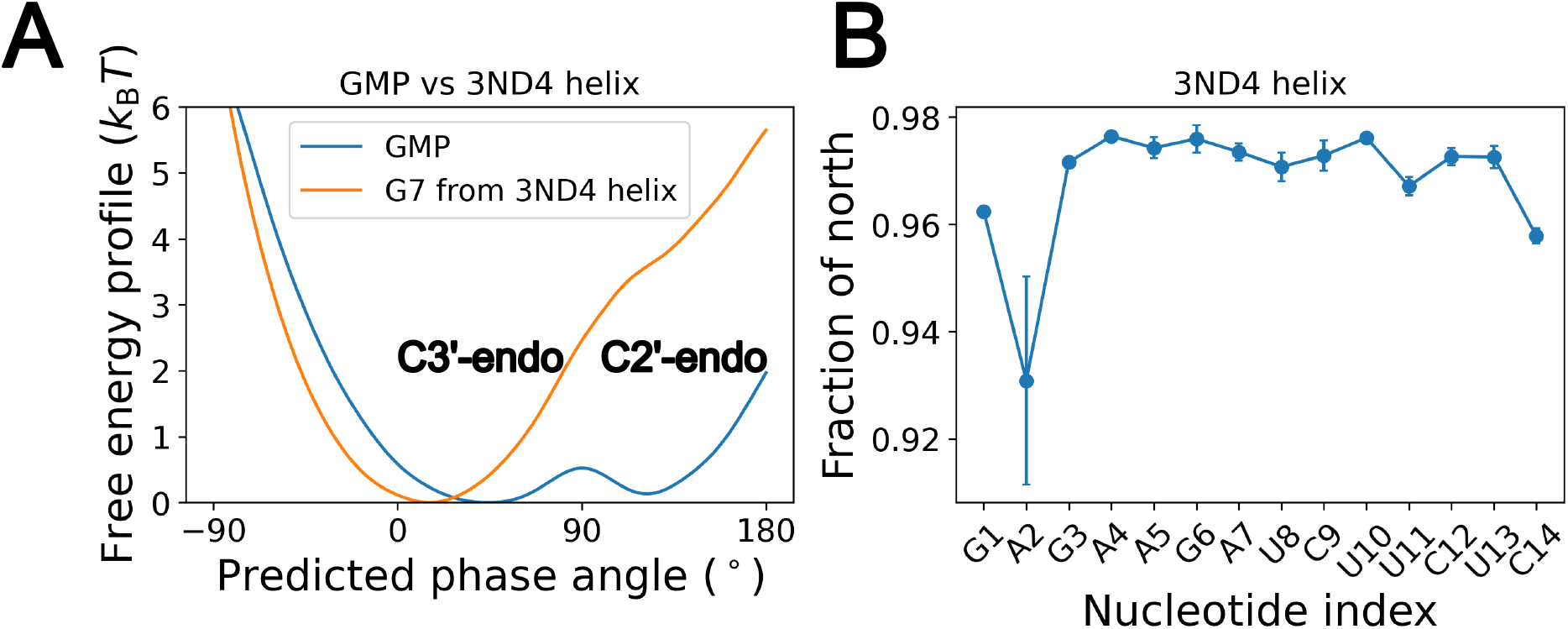
Sugar puckering FES and fractions of north conformations. (A) FES of the predicted phase angle of a guanosine monophosphate (GMP) and the G7 in a 3ND4 helix. (Negative phase angles (−90° − 0°) are shifted from the range (270° − 360°)) (B) Equilibrium fraction of north conformation of each nucleotide over the nucleotide indices. (The north population is defined as having a phase angle in the range 0° − 90° or 270° − 360°, which are usually dominated by C3’-endo, see definitions in the section Sugar Puckering Energy Function and Fig. 2A). Error bars are calculated using the last 3/4 of the trajectory. The terminal nucleotides A0 and U15 are not included.

For G7 in the A-form helix (PDB 3ND4), the local minimum of C2’-endo disappears and becomes a shoulder with free energy 4–5 *k*_B_*T* higher than the C3’-endo (Fig. 8A). Along the double helix, CRANBERRY predicts that the fraction of north conformations remains at 94–96% (Fig. 8B). (“North” includes C3’-endo and nearby states; we report north fractions to capture contributions from conformations near C3’-endo, e.g., C2’-exo, C4’-exo.) We did not tune specifically for this behavior; rather, it emerges for the nucleotides in the double helix after the ConDiv training and fine-tuning. The resulting strong preference for north puckers is qualitatively consistent with their predominance in A-form RNA helices and with the occurrence of C2’-endo primarily in nonhelical regions, including bulges, loops, and termini (81–83, 89, 126, 127). Using an all-atom force field, the free energy difference Δ*G*_3↔2_ is predicted to be about +10 *k*_B_*T* and the barrier is estimated to be about 20 *k*_B_*T* (83). The qualitative agreement between CRANBERRY and the all-atom simulations can be explained by the following mechanisms for C2’-endo destabilization in A-form helices.

The C2’-endo state may be disfavored in A-form helices through several coupled mechanisms. C2’-endo puckering can disrupt base stacking (83), an effect represented in CRANBERRY through its sugar puckering and base-stacking potentials. Steric interactions involving the 2’-OH and hydration effects may also shift the conformational equilibrium toward C3’-endo (81, 83, 128). These effects are not yet fully captured by CRANBERRY, and may contribute to its quantitative difference from the all-atom predictions (83). However, to the best of our knowledge, no direct experimental measurements exist for the free-energy difference or the height of the barrier.

## Limitations

Similarly to other CG models (53, 62), CRANBERRY treats electrostatics at the level of the Debye-Hückel approximation, which is strictly valid only at low salt concentrations. Although CRANBERRY captures salt-dependent size changes in disordered RNA, its accuracy for ion-dependent conformational ensembles at higher salt concentrations remains unclear. In the future, explicit ion models may need to be developed, as demonstrated in other CG models (5, 60, 129).

Another limitation is that CRANBERRY currently cannot model divalent ions such as magnesium or base-ion interactions. The only electrostatic interactions currently explicitly included are repulsions between the phosphates. CRANBERRY is not expected to model ion-dependent structures such as G-quadruplexes. In the future, CRANBERRY could be extended to include these terms.

CRANBERRY can qualitatively capture sugar puckering. In helical regions, CRANBERRY does not destabilize south conformations (or mostly, C2’-endo) enough compared to all-atom models. This is primarily due to the absence of explicit 2’-OH and backbone sterics in the coarse-grained model. This effect could be addressed by coupling the *η* − *θ* map with sugar puckering, favoring the C3’-endo in the helical regimes of the *η* − *θ* map (86, 130, 131). There are also known correlations between the map *η* − *θ* and sugar puckering states (86), which could be included as well. However, these are considered second-order effects and are omitted in our initial CRANBERRY implementation. Introducing the *η* − *θ* map might also help stabilize various turn motifs in RNA.

CRANBERRY currently models only the 25 most frequent base-pairing geometries, which together account for 87% of cases in the fragment data set (Materials and Methods: Data sets). We note that our approaches are entirely compatible with new experimental data or more accurate *ab initio* calculations. In the future, all 156 known base-pairing geometries could be parameterized and incorporated. Also, current native stability data suggest that the noncanonical base-pairing interactions could be strengthened or rebalanced. Future parameterization could incorporate quantum-mechanical interaction-energy surfaces and experimentally inferred opening free energies for individual base pairs and tertiary contacts, such as those accessible through recently developed quantitative chemical-probing approaches (132).

CRANBERRY currently lacks an atomistic backmapping procedure, limiting direct comparison of its structural ensembles with atom-resolved experimental observables. Developing such a procedure, together with calibration of internal dynamical timescales, would enable future validation against NMR-derived conformational ensembles and relaxation data.

## Conclusions

We highlight the following design elements that enable CRANBERRY’s performance: contrastive divergence training (71), the fine-tuning process that greatly improves the disordered states and folding cooperativity, and the incorporation of physically-motivated terms such as non-canonical base pairing and sugar puckering.

CRANBERRY’s major accomplishments are:

1. Stabilizes helices, tetraloops, and pseudoknots at room temperature within 2–6 Å RMSD of the crystal/NMR structures;
2. Captures the measured NaCl-concentration dependence of disordered rU30 size and reproduces its SAXS profiles across the tested concentration range;
3. Reproduces the scale and sequence dependence of the dinucleotide stacking free energies used as a fine-tuning target;
4. Recovers cooperative melting of the tetraloop ggcGCAAgcc and double helix cacag within 15 K of experiments but exhibits better cooperativity compared to all-atom;
5. Folds ggcGCAAgcc *de novo*, reversibly from an extended strand with minimum RMSD as 1.4 Å;
6. Reproduces sugar-pucker preferences in mononucleotides and helices.

We developed CRANBERRY, a six-site coarse-grained model for RNA that simultaneously captures sugar-pucker transitions, noncanonical base pairing, native and disordered states, and thermodynamics. Employing a linear-regression-based many-body bonded potential for sugar puckering transitions, CRANBERRY naturally reproduces the equilibrium populations of ribose puckers. Our contrastive-divergence training, augmented with fine-tuning against experimental stacking free energies and disordered-RNA Rgs, produces nonbonded potentials that balance native-state stabilities with realistic melting behavior. CRANBERRY’s computational efficiency opens the door to routine exploration of RNA folding landscapes. We anticipate that CRANBERRY will serve as a practical tool for simulating complex RNA assemblies.

## ACKNOWLEDGEMENTS

The authors thank Stefano Piana, Jannette Carey, Kathleen Hall, Modesto Orozco, and Lois Pollack for insightful discussions. The authors also acknowledge the computational resources provided by the Research Computing Center at the University of Chicago. ChatGPT (OpenAI) was used for language editing and post hoc optimization of code in the GitHub repository. All AI-assisted code changes were reviewed and tested by the authors, who take full responsibility for the manuscript and deposited code.

## Data availability

The CRANBERRY source code, force-field parameters, simulation examples, and analysis scripts used in this study are available at https://github.com/depablogroup/cranberry.

## Author contributions

Y.W.: Conceptualization, Methodology, Software, Formal Analysis, Investigation, Visualization, Writing–Original Draft, and Writing–Review & Editing. R.A.: Conceptualization, Methodology, Investigation, Formal Analysis, and Writing–Review & Editing. A.E.C.: Conceptualization and Writing–Review & Editing. X.P.: Conceptualization and Writing–Review & Editing. P.F.Z.R.: Investigation and Writing–Review & Editing. K.L.: Methodology and Writing–Review & Editing. C. S.: Investigation and Formal Analysis. K.T.: Investigation and Formal Analysis. T.R.S.: Conceptualization, Supervision, Funding Acquisition, and Writing–Review & Editing. J.J.d.P.: Conceptualization, Supervision, Funding Acquisition, and Writing–Review & Editing.

